# Nuclear 2′-O-methylation regulates RNA splicing through its binding protein FUBP1

**DOI:** 10.1101/2025.04.20.649728

**Authors:** Boyang Gao, Bochen Jiang, Zhongyu Zou, Bei Liu, Weijin Liu, Li Chen, Lisheng Zhang, Chuan He

**Affiliations:** Department of Molecular Genetics and Cell Biology, University of Chicago, Chicago, IL 60637, USA; Department of Chemistry, The University of Chicago, Chicago, IL 60637, USA; Howard Hughes Medical Institute, The University of Chicago, Chicago, IL 60637, USA; Division of Life Science, The Hong Kong University of Science and Technology (HKUST), Kowloon, Hong Kong SAR, China; Department of Chemistry, The Hong Kong University of Science and Technology (HKUST), Hong Kong SAR, China

## Abstract

2′-O-methylation (N_m_) is an abundant RNA modification exists on different mammalian RNA species. However, potential N_m_ recognition by proteins has not been extensively explored. Here, we employed RNA affinity purification followed by mass spectrometry to identify N_m_-binding proteins. The candidates exhibit enriched binding at known N_m_ sites. Interestingly, some candidates display nuclear localization and functions. We focused on the splicing factor FUBP1. Electrophoretic mobility shift assay (EMSA) validated preference of FUBP1 to N_m_-modified RNA. As FUBP1 predominantly binds intronic regions, we profiled N_m_ sites in chromatin-associated RNA (caRNA) and found N_m_ enrichment within introns. Depletion of N_m_ led to increased exon skipping, suggesting N_m_-dependent splicing regulation. The caRNA N_m_ sites overlap with FUBP1 binding sites, and N_m_ depletion reduced FUBP1 occupancy on modified regions. Furthermore, FUBP1 depletion induced exon skipping in N_m_-modified genes, supporting its role in mediating N_m_-dependent splicing regulation. Overall, our findings identify FUBP1 as an N_m_-binding protein and uncover previously unrecognized nuclear functions for RNA N_m_ modification.

## Introduction

Throughout the life cycles of RNA, chemical modifications play critical roles in regulating RNA processing, metabolism and function(*1, 2*). These modifications can alter the physical properties of the RNA molecule(*3, 4*), or recruits specific binding proteins (readers) to modulate RNA function and subsequent cellular pathways(*5–9*). While changes in physical properties primarily influence RNA structure-dependent regulation, the recruitment of reader proteins can impact diverse downstream processes, including splicing(*10*), degradation(*5*), translation(*6, 9*), and RNA transport to specialized cellular locations(*11–13*). Additionally, recruitment of binding proteins may alter the surrounding state of the modified RNA; examples include chromatin state regulation by N^6^-methyladenosine methylation of chromatin-associated RNA (caRNA)(*7, 8, 14*).

N_m_ is one of the most abundant modifications(*15*). It can be found in almost all RNA species, including ribosomal RNA (rRNA), transfer RNA (tRNA), small nuclear RNA (snRNA), small nucleolar RNA (snoRNA), microRNA (miRNA) and messenger RNA (mRNA). It has been shown to regulate ribosome biogenesis(*16, 17*), gene expression(*18, 19*), innate immune sensing(*20, 21*) and cell fate decisions(*22*). N_m_ entails methylation at the 2′-OH position of the ribose on the RNA backbone. Consequently, it can occur on any of the four ribonucleotide residues, namely A_m_, G_m_, C_m_ and U_m_. The methylation further stabilizes the ribose 3′-endo conformation, favoring the A-type RNA helix and restricting strand flexibility(*23–25*). Consequently, N_m_ installation alters physical properties of the modified RNA. This effect orchestrates N_m_-dependent cellular functions of different RNA species, such as stabilizing ribosome structure through modified rRNA(*26*), and controlling translational efficiency through internal mRNA N_m_ modification(*18*).

However, whether N_m_ could be recognized by binding proteins (readers) remains largely unexplored, particularly within the internal regions of mRNA or pre-mature mRNA. While previous studies have identified over a thousand internal N_m_ sites(*27*), the corresponding function remains elusive. We speculated that N_m_-binding proteins may exist to regulate processing, metabolism or function of modified RNA.

In this work, we conducted RNA affinity purification followed by mass spectrometry (MS) to identify candidate N_m_-binding proteins, among which we focused on the splicing factor FUBP1. Upregulation of FUBP1 has been suggested to promote proliferation of multiple types of cancers(*28, 29*). Initially characterized as a transcription factor modulating MYC expression, FUBP1 is now recognized as an RNA-binding protein (RBP) involved in pre-mRNA splicing(*30, 31*). Previous studies have indicated that FUBP1 stabilizes splicing machineries at 3′ splice sites, including U2AF2 and SF1.

Additionally, it interacts with components of U1 snRNPs, potentially facilitating splice sites pairing in long introns. Understanding the binding specificity of FUBP1 could elucidate the mechanism underlying alternative splicing (AS) events which could be affected by N_m_ modification through FUBP1.

Using electrophoretic mobility shift assay (EMSA), we confirmed the binding preference of FUBP1 to N_m_-modified oligos, supporting its role as an N_m_-binding protein. N_m_-mut-seq(*27*) analysis of caRNA in HepG2 cells identified 5,575 N_m_ sites, with more than half of the intragenic N_m_ sites localized in introns. Disruption of N_m_ installation led to altered splicing patterns, especially increased exon skipping. These N_m_ sites were bound by FUBP1 in a manner responsive to N_m_ depletion. Finally, FUBP1 preferentially regulates exon skipping at N_m_-modified regions. Taken together, our study identifies FUBP1 as, to our knowledge, the first example of an N_m_-binding protein and highlights a previously unrecognized role of N_m_ in splicing regulation.

## Results

### RNA affinity purification followed by MS identified candidate N_m_-binding proteins

To identify proteins that may preferentially recognize internal RNA N_m_ modification, we designed biotinylated RNA probes based on published internal N_m_ sites in HepG2(*27*). Given that N_m_ can occur on four distinct types of ribonucleotides (A_m_, G_m_, C_m_ and U_m_), probes with a single type of N_m_ may not fully capture the binding proteins recognizing four different N_m_ modifications. Considering the predominant presence of G_m_ and A_m_ in mRNA N_m_ modifications(*27*), we designed two probes: one with G_m_ modification (G_m_-1) and the other with A_m_ modification (A_m_-2) (Fig. 1A). Each probe was accompanied by a corresponding control oligo lacking the N_m_ modification, denoted as Ctrl-1 and Ctrl-2.

**Fig. 1.**
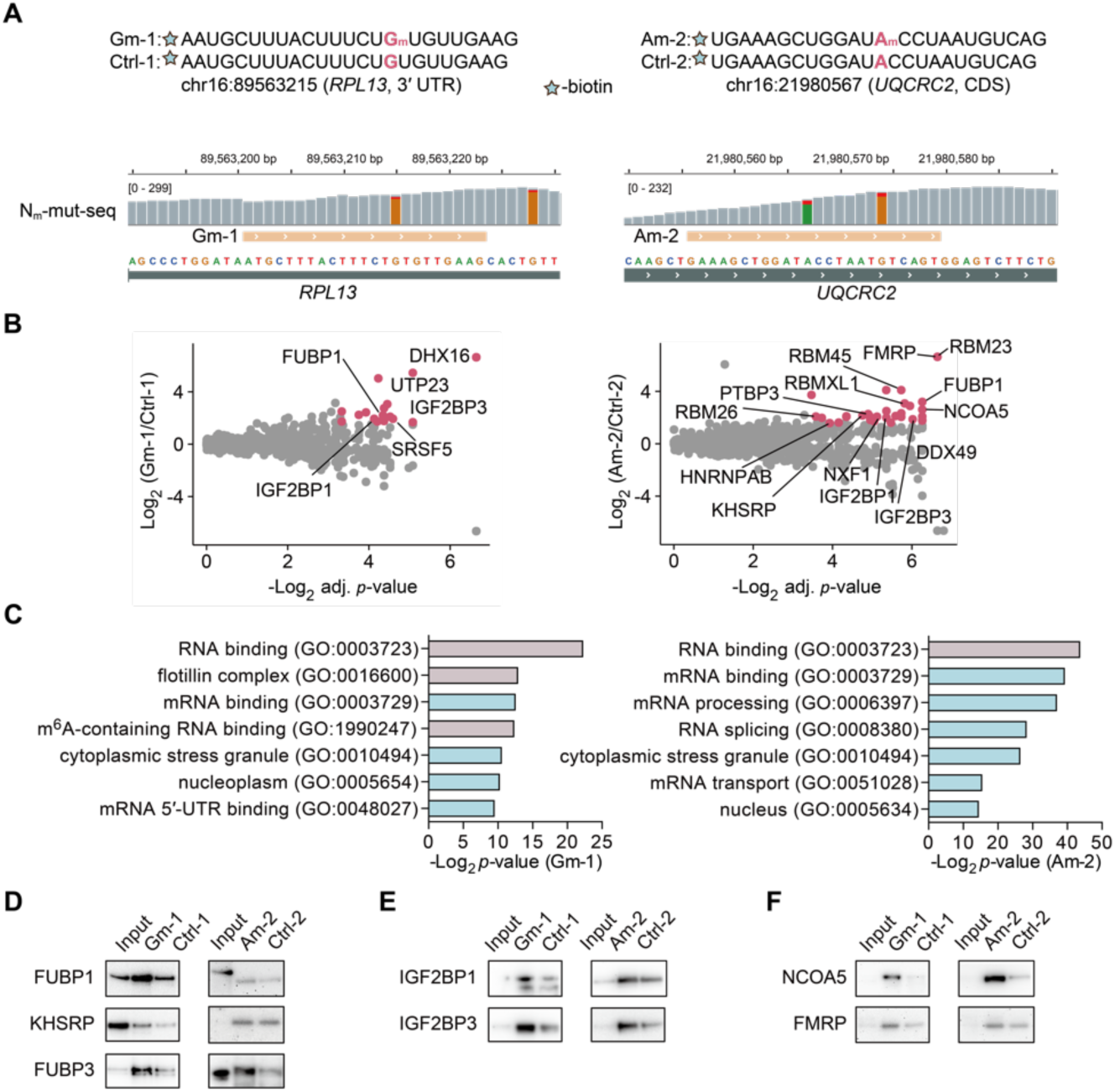
RNA affinity purification followed by proteomics identified candidate N_m_ binding proteins. (**A**) Design of Gm-1 and Am-2 probes based on known N_m_ sites in HepG2 mRNA. Integrative genomics viewer (IGV) tracks of published N_m_-mut-seq showed read depth of mutated (red) and un-mutated reads (G in orange, A in green). (**B**) Peptides from proteins that may preferentially bind to RNA Gm-1 (left) and Am-2 (right) probes over control unmodified probes. The measured peptide abundance was normalized to the total peptide amount, and the mean of the peptide group abundance was adjusted to the same for different samples. The geometric median of 3 replicates was used for fold change calculation between N_m_-modified and unmodified samples. N_m_-binding protein candidates (red) should have log_2_ fold change > 1.58, adjusted *p*-value < 0.1 and peptide number ≥ 4. (**C**) GO analysis of candidate N_m_-binding proteins. (**D**) Immunoblotting of candidate N_m_-binding protein enrichment by oligo pull-down. The ratio of the corresponding lysate amount of input vs IP samples are 1:100.

These probes were selected based on two reported N_m_ sites, with relatively high mutation ratio and responsiveness to the knockdown (KD) of their methyltransferase FBL(*27*) (Fig. 1). The Gm-1 sequence is in the 3′-untranslated region (3′ UTR) of *RPL13*, while the Am-2 sequence situates in the coding sequence (CDS) of *UQCRC2*.

We performed affinity purification with Gm-1 and Am-2 using HepG2 cell lysate, respectively, and employed MS for protein identification of the pulldown fraction. Peptide abundance was normalized to the total peptide abundance, and fold changes of N_m_-modified versus control groups were computed. With a cutoff of log_2_ fold change > 1.58, adjusted *p*-value < 0.1 and peptide number ≥ 4, we identified 44 enriched proteins in the Gm-1 group and 38 enriched proteins in Am-2 group (Fig. 1B, Fig. S1A). These enriched proteins are candidate binding proteins for G_m_ and A_m_ modifications. The identified protein candidates include mRNA binding proteins and proteins involved in RNA-related metabolic pathways, based on gene ontology (GO) analysis (Fig. 1C). Interestingly, GO terms associated with the nucleus functions are enriched in both Gm-1 and Am-2 enriched protein candidates. This suggests nuclear functions of N_m_ modifications that has not be previously recognized.

In the candidate protein list, we observed proteins present in both G_m_ and A_m_ enriched groups, indicating them as potential N_m_ binders without base specificity. One example is FUBP1; interestingly, two other homologs within the same protein family, FUBP3 and KHSRP, were also identified in the A_m_ candidate list. Although their enrichment in the Gm-1/Ctrl-1 sample pair did not surpass the cutoff threshold, the structural similarity within the protein family suggests all three proteins may preferentially bind to N_m_, as the lack of enrichment in the Gm-1/Ctrl-1 sample pair could be due to insufficient coverage of the peptide pools. To validate this speculation, we detected the protein enrichment by immunoblotting after RNA affinity purification using the two pairs of probes (Fig. 1A). As expected, both FUBP1 and FUBP3 were enriched by the two N_m_ probes compared to their respective controls (Fig. 1D). KHSRP showed enrichment in Gm-1/Ctrl-1 sample pair, but its enrichment in the Am-2/Ctrl-2 sample pair was minimal. This observation indicates that FUBP1 and FUBP3 are robust candidates recognizing both G_m_ and A_m_ modifications on RNA.

Additionally, IGF2BP1 and IGF2BP3 emerged as candidate binders of both G_m_ and A_m_. IGF2BPs were initially identified as m^6^A-binding proteins that stabilize target mRNAs and facilitates their storage under stress conditions(*32*). Our previous work also revealed their role in recognizing m^7^G(*33*). While IGF2BP1 primarily binds m^6^A and stabilizes associated mRNAs, IGF2BP3 preferentially recognizes m^7^G and promotes mRNA degradation. Immunoblotting following affinity purification of both probe pairs confirmed the preferentially binding of IGF2BP1 and IGF2BP3 to N_m_-modified RNA (Fig. 1E). These findings may suggest a complex mechanism of RNA modification recognition by the IGF2BP proteins.

In addition to the FUBP and IGF2BP RBP families, we also validated enrichment of FMRP and NCOA5. FMRP and NCOA5 were enriched by both Gm-1/Ctrl-1 and Am-2/Ctrl-2 probe pairs through affinity purification followed by immunoblotting (Fig. 1F). Notably, NCOA5 is a transcriptional coactivator that interacts with estrogen receptors(*34*). We analyzed a published RNA-binding region identification (RBR-ID) dataset(*35*) and observed enriched NCOA5 peptides, indicating its involvement in RNA binding as an RBP (Fig. S1B). NCOA5 could be another example of a transcription factor that binds to and is potentially regulated by RNA. Its preferential binding to N_m_ may suggest a possible role for N_m_ in transcriptional regulation. Overall, we validated 7 proteins with binding preferences for N_m_-modified probes: FUBP1, KHSRP, FUBP3, IGF2BP1, IGF2BP3, FMRP and NCOA5.

### Candidate N_m_-binding proteins are enriched at internal mRNA N_m_ sites

To provide further cellular evidence that these proteins bind N_m_ modifications, we investigated their RNA binding profiles using publicly available eCLIP datasets from the ENCODE project(*36, 37*). Among the seven identified N_m_ binders, KHSRP, FUBP3, IGF2BP1 and IGF2BP3 have eCLIP data generated in HepG2 cells. Metagene plots generated from these datasets revealed enriched binding of all four proteins around the reported confident N_m_ sites (Fig. 2A-D)(*27*), supporting the preferences of these proteins towards N_m_ modification.

**Fig. 2.**
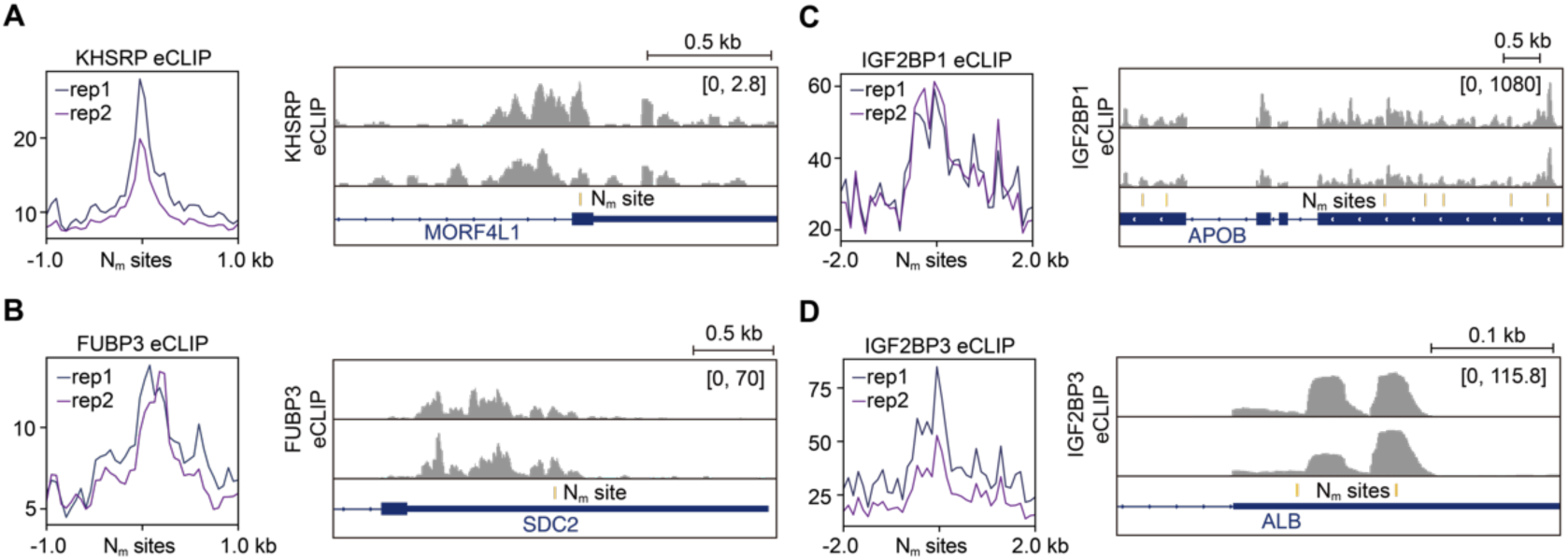
RNA-binding sites of candidate N_m_-binding proteins were enriched at N_m_ sites. (**A-D**) Metagene plots of eCLIP signals at HepG2 mRNA N_m_ sites (left) and representative IGV tracks (right) from binding sites of candidate N_m_ binding proteins KHSRP (A), FUBP3 (B), IGF2BP1 (C), and IGF2BP3 (D).

We examined the sequence contexts of all four proteins at their N_m_ binding sites. Their enriched N_m_ motifs closely resembled their own RNA binding motifs (Fig. S2A-D), suggesting that the binding selectivities of these N_m_-binding proteins are determined by their canonical sequence contexts but likely further enhanced by the N_m_ modification. To further investigate how RNA-binding preferences affect N_m_ site selectivity, we analyzed the distribution of their binding targets and associated N_m_ sites across mRNA regions. While N_m_ sites were generally enriched in the CDS compared to the 3’UTR and 5’-untranslated regions (5’UTR), N_m_-binding proteins with 3’UTR preferences, KHSRP and FUBP3, predominantly bound to N_m_ sites within the 3’UTR. This highlights the preferences of these RBPs to their canonical binding sites. Conversely, IGF2BP1 and IGF2BP3 exhibited a stronger enrichment of bound N_m_ sites in the CDS than their overall binding sites, suggesting a potentially important contribution of selective N_m_ installation to RNA binding by these RBPs.

The reported function of IGF2BP1 in stabilizing mRNA aligns with the general effect of N_m_ on mRNA levels as previously described(*19, 27, 38*). Accordingly, we analyzed RNA level changes using published KD datasets for *IGF2BP1* in HepG2 cells(*32*). N_m_-modified mRNA transcripts showed a more notable decrease in expression following *IGF2BP1* KD compared to unmodified transcripts (Fig. S2F), supporting the preferential interaction of IGF2BP1 with N_m_ modifications. Further investigation into IGF2BP1 binding to methylated RNA is needed to elucidate the interplay among m^6^A, m^7^G and N_m_ modifications.

### FUBP1 preferentially binds internal mRNA N_m_ sites

The enrichment of nuclear proteins in the N_m_ binder candidate list suggests intriguing functions of N_m_ in the cell nucleus. We chose FUBP1 as an example for more detailed investigations, as FUBP1 is a well-characterized splicing factor(*28, 31*). Its preference for N_m_ has been validated by using the two probe pairs in both proteomics and immunoblotting analyses. Nevertheless, the RNA affinity purification approach leaves a possibility of indirect FUBP1 binding mediated by an N_m_-recognizing partner protein. To confirm the direct interaction between FUBP1 and the N_m_ modification, we recombinantly expressed and purified FUBP1 with a C-terminal strep-tag^®^ from the Expi293 expression system (Fig. S3A). We then assessed the binding affinity of the purified FUBP1 towards two probe pairs by electrophoretic mobility shift assay (EMSA). We observed preferential binding of FUBP1 to both N_m_-modified RNA probes compared to their respective controls (Fig. 3A). As a sugar 2’-OH modification, N_m_ recognition by FUBP1 could be more subtle, when compared with other well-recognized RNA modifications such as m^6^A by the reader YTHDF proteins(*39*). Our EMSA results support that the preferential binding of FUBP1 to N_m_ is mediated by a direct interaction. Future structural characterization may reveal details of this recognition.

**Fig. 3.**
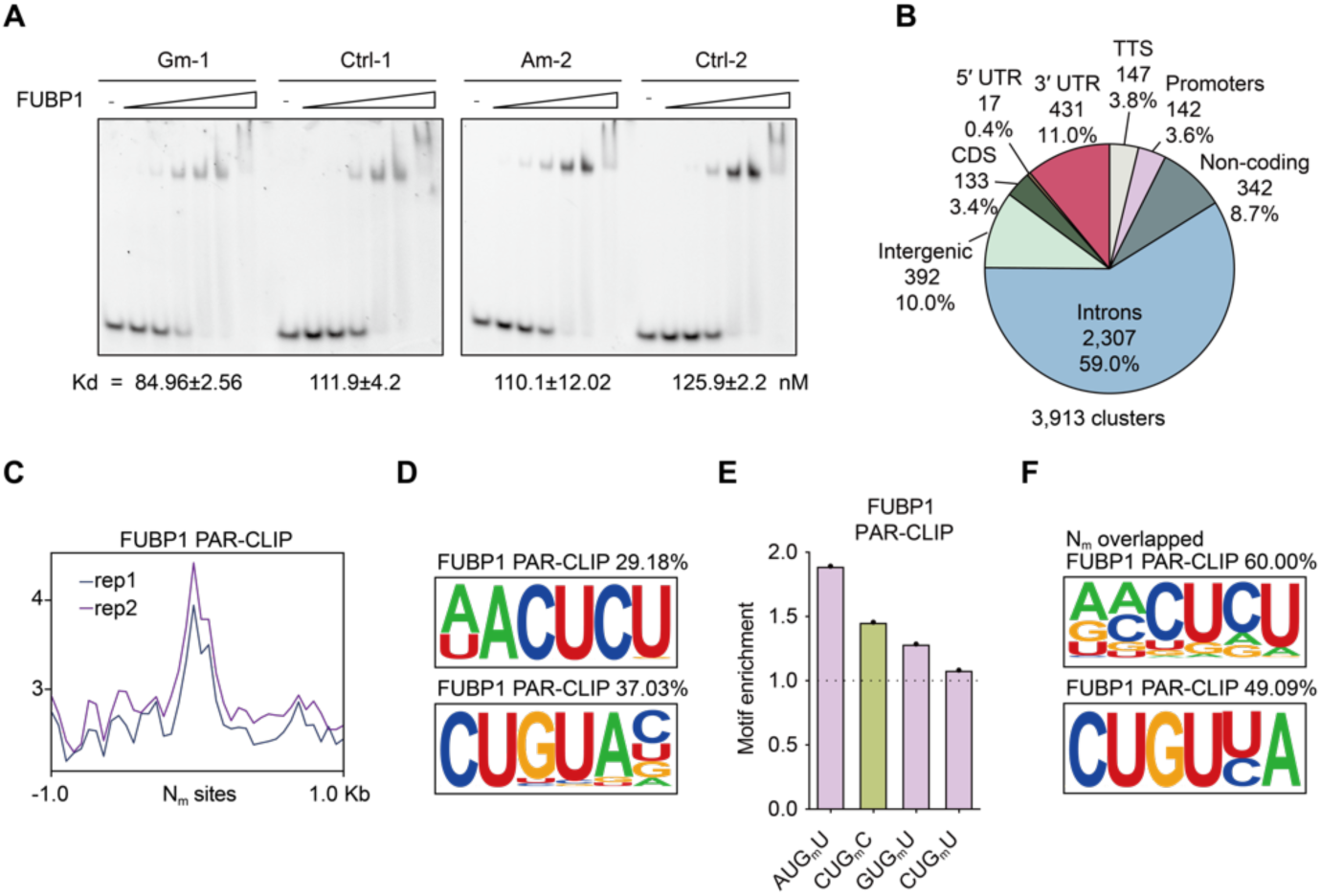
Splicing regulator FUBP1 prefers N_m_-modified RNA. (**A**) EMSA of FUBP1 binding towards Gm-1 and Am-2 probes versus their controls. Probe concentration: 20 nM. Protein concentration: starting from 400 nM with 2-fold dilution. (**B**) Distribution of FUBP1 PAR-CLIP binding clusters across transcript elements. TTS: transcription termination sites. (**C**) Metagene plot of FUBP1 PAR-CLIP signals at HepG2 mRNA N_m_ sites. (**D**) FUBP1 binding motifs identified in PAR-CLIP. (**E**) Enrichment of N_m_ motifs at the FUBP1 binding sites. N_m_ motifs resembling the FUBP1 RNA binding motifs are colored in pink. N_m_ motifs that are also enriched by KHSRP or FUBP3 binding are colored in green. (**F**) FUBP1 binding motifs identified in PAR-CLIP clusters overlapping with mRNA N_m_ sites.

In addition to biochemical evidence, we examined whether FUBP1 binding occurs at endogenous N_m_ sites within various RNA sequence contexts. We conducted Photoactivatable Ribonucleoside-Enhanced Crosslinking and Immunoprecipitation (PAR-CLIP) of FUBP1 in HepG2 cells. Genome-wide mutation analysis revealed T-to-C mutation ratio > 55% (Fig. S3B). We identified 3,913 FUBP1 binding sites (Fig. S3C), with ∼59% of these located within intronic regions (Fig. 3B), consistent with the reported intron preference of FUBP1 binding(*31*). With the published mRNA N_m_ sites at base-resolution, we first investigated FUBP1 binding at reported mature mRNA N_m_ sites. A metagene plot revealed a clear enrichment of PAR-CLIP reads at the confident N_m_ sites (Fig. 3C). FUBP1 binding motifs were found to be U-rich (Fig. 3D), consistent with the U-rich motifs associated with mRNA N_m_. Notably, N_m_ motifs enriched at FUBP1-binding sites closely resembled the RNA-binding motifs shared across all its binding sites (Fig. 3E-F). These results indicate an intrinsic preference of FUBP1 for N_m_-modified regions, further supporting its role as an N_m_-binding protein.

### Abundant N_m_ modifications on caRNA regulates splicing events

The preference of splice factor FUBP1 for N_m_ modifications indicates an unrecognized intron-dependent function of N_m_. To explore whether N_m_ modifications are also present in intronic regions, we measured the overall levels of N_m_ in rRNA-depleted HepG2 caRNA by Ultrahigh-Performance Liquid Chromatography coupled with triple Quadrupole Mass Spectrometry (UHPLC-QQQ-MS/MS). Intriguingly, the intensity ratios of A_m_/A and C_m_/C are significantly higher in caRNA compared to mRNA (Fig. 4A). The G_m_/G ratio in caRNA is about one third of that in mRNA; however, the considerably longer length of introns likely compensates, resulting in an overall more G_m_ sites in introns than those in exons. Our findings suggest enrichment of N_m_ modifications in caRNA.

**Fig. 4.**
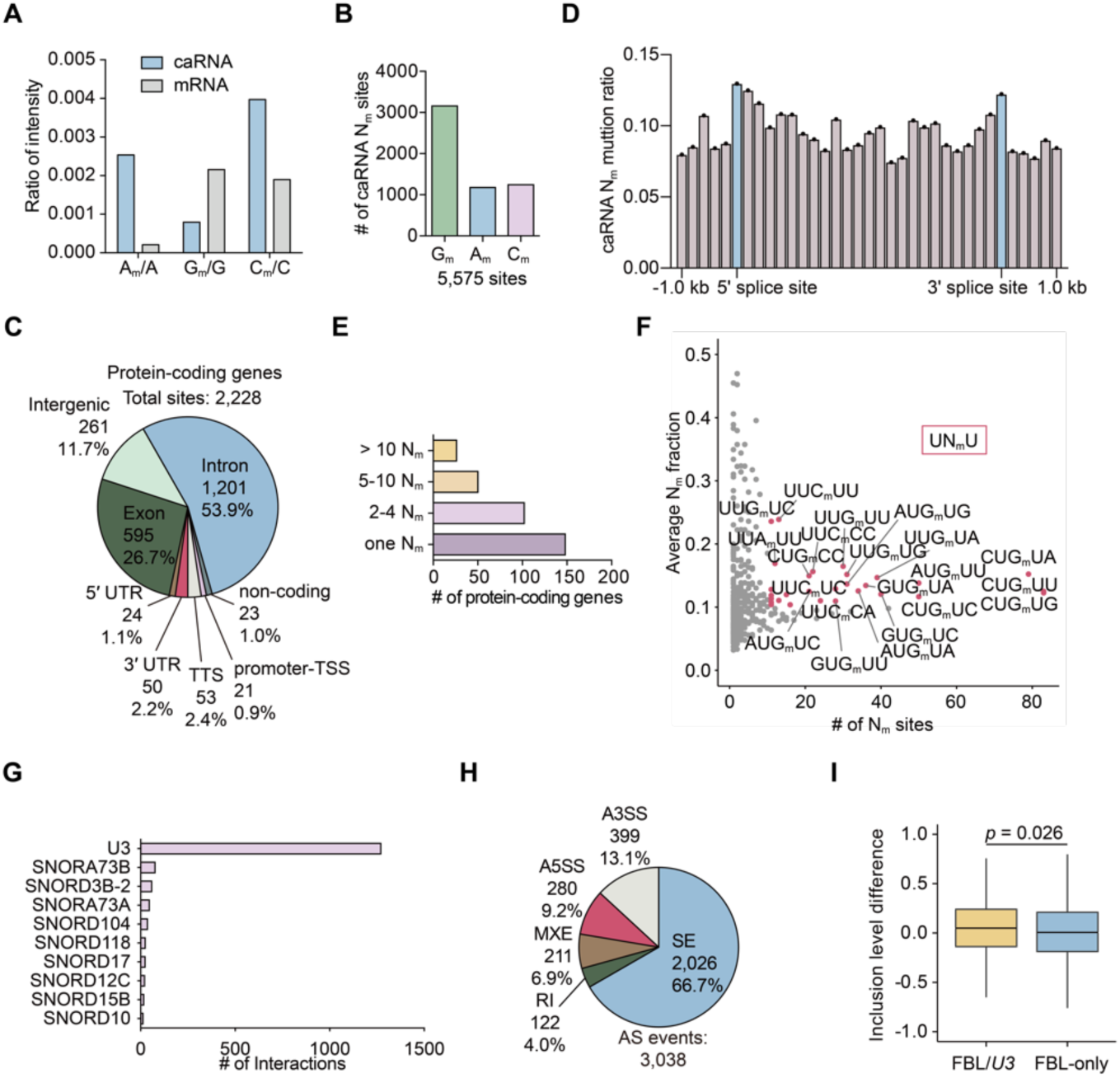
N_m_ modifications on HepG2 caRNA affect splicing events. (**A**) UHPLC-QQQ-MS/MS of N_m_ abundances in HepG2 mRNA and caRNA. (**B**) Composition of G_m_, A_m_, and C_m_ among the 5,575 caRNA N_m_ sites. (**C**) Transcript element distribution of 2,228 caRNA N_m_ sites in protein-coding genes. (**D**) Averaged caRNA N_m_ mutation ratio across intronic regions in a 200-bp binning. Intronic sequences at the boundary of splice sites were marked in blue. (**E**) Profile of modified protein-coding gene transcripts with the corresponding numbers of N_m_ sites. (**F**) Number and average mutation ratio of all 5-base N_m_ motifs. UN_m_U was enriched in more frequent and highly modified motifs. (**G**) Number of interactions with various snoRNA detected by snoKARR-seq at all caRNA N_m_ sites. (**H**) Profile of FBL-dependent AS events, with significance threshold FDR < 0.1. (**I**) Inclusion level differences of FBL/*U3* or FBL-only SE events.

To examine caRNA N_m_ distribution profiles, we performed N_m_-mut-seq on HepG2 caRNA. N_m_-mut-seq employs an imbalanced dNTP supply in reverse transcription (RT) reactions to induce mutations of G_m_, A_m_ and C_m_ into T(*27*). During analysis, the treated samples could be controlled by RT reactions using balanced dNTP supply (input), as well as a set of spike-in that does not harbor N_m_ modifications but undergoes RT with imbalanced dNTP (background). We analyzed the mutations of caRNA N_m_-mut-seq by JACUSA(*40*), and designated sites as N_m_-modified when all three replicates showed sequencing depth ≥ 10, mutated read depth ≥ 3, mutation ratios > 3-fold of input mutation ratios, and > 1.5-fold of background mutation ratios. Based on these criteria, we identified 5,575 N_m_ sites, far more than the reported 1,051 N_m_ sites or 494 confident N_m_ sites found in HepG2 mRNA(*27*).

Similar to mRNA, G_m_ sites are the most abundantly modified N_m_ sites in caRNA (Fig. 4B), despite the relatively higher abundances of A_m_ and C_m_ detected by UHPLC-QQQ-MS/MS (Fig. 4A). The numbers of confident A_m_ and C_m_ sites in caRNA are comparable. The majority of caRNA N_m_ sites in caRNA are located within protein-coding genes (Fig. S4A), with ∼53.9% of them residing in introns (Fig. 4C). This is consistent with our hypothesis of intron-dependent function of N_m_. These intronic N_m_ sites show modest enrichment near 5′ and 3′ splice sites (Fig. 4D), suggesting a potential role of N_m_ in splicing regulation. Our previous work has shown that internal N_m_ sites on mRNA often appear densely clustered along a stretch of RNA(*38*). We hypothesized that such clustering may amplify the relatively weak binding preferences of N_m_-binding proteins toward modified RNA. Indeed, we observed that 2,228 caRNA N_m_ sites on protein coding genes were distributed across only 323 genes, with more than half of these genes harboring multiple N_m_ sites (Fig. 4E). In addition to protein-coding genes, several long non-coding RNAs (lncRNAs) also accumulate hundreds of N_m_ sites (Fig. S4B), suggesting a previously unrecognized layer of N_m_-dependent regulation in their respective functions.

N_m_ installation in HepG2 caRNA is enriched in the UN_m_U sequence context (Fig. 4F), similar to the motifs observed in HepG2 mRNA(*27*). The consistency should be expected, as caRNA modifications in intronic regions and non-coding RNA (ncRNA) are generally installed by the same methyltransferases responsible for mRNA modification, despite their different fates after RNA processing and nuclear export(*41*). The observed motifs also overlap with U-rich elements recognized by key splicing factors, such as the 5’ splice sites GU, the branch sites, the UGCAUG hexanucleotides(*42*), and the polypyrimidine tract(*43*), *etc.* This further hints a role of N_m_ in splicing regulation. Notably, the U-rich motif is consistent with the known RNA-binding preference of FUBP1(*31*), further suggesting its role as an N_m_-binding protein that recognizes intronic N_m_ to potentially modulate splicing.

To understand the regulation of N_m_ installation on caRNA, we analyzed snoRNA interactions with N_m_-modified regions using the published snoKARR-seq dataset(*44*). The majority of snoRNA interaction with caRNA N_m_ sites were mediated through *U3* (Fig. 4G and S4C), highlighting the critical role of *U3* in directing N_m_ installation on caRNA. Our finding suggests that *U3* KD may serve as an effective strategy to perturb caRNA N_m_ installation in addition to depletion of the N_m_ methyltransferase FBL.

To investigate the role of caRNA N_m_ in splicing regulation, we depleted FBL or *U3* in HepG2 cells (Fig. S4D). We examined five types of AS events: alternative 3’ splice sites (A3SS), alternative 5’ splice sites (A5SS), mutually exclusive exons (MXE), retained introns (RI), and skipped exons (SE). AS events dependent on FBL or *U3* exhibited similar distributions, with SE events being the most prevalent (Fig. 4H and S4E). In both FBL depletion and *U3* KD, more SE events were upregulated than downregulated, while other AS types showed non-significant or inconsistent changes (Fig. S4F). This indicates that N_m_ installation may play a role in preventing exon skipping. To further examine this, we compared SE events that consistently occurred following both *FBL* and *U3* KD (FBL/*U3*) with those that occurred only or inconsistently after *FBL* KD (FBL-only) or *U3* KD (U3-only). The FBL/*U3* SE events exhibited a higher degree of upregulation compared to FBL-only and *U3*-only groups (Fig. 4I and S4G), further supporting a role for N_m_ in suppressing SE events. Overall, our findings reveal abundant N_m_ modifications in caRNA, with intronic installation affecting splicing regulation.

### FUBP1 mediates N_m_-dependent splicing regulation

Our N_m_-mut-seq analysis of HepG2 caRNA revealed a substantial presence of N_m_ modifications within intronic regions, potentially involved in splicing regulation. Given that FUBP1 is a known splicing factor implicated in splice site recognition(*31*), we hypothesized that it may mediate N_m_-dependent splicing regulation. To investigate this, we first validated the association of FUBP1 with icaRNA N_m_ sites. Of the 5,575 caRNA N_m_ sites, 2,030 were bound by FUBP1, as determined by PAR-CLIP analysis (Fig. 5A). Correspondingly, FUBP1 PAR-CLIP signals were enriched around caRNA N_m_ sites (Fig. 5B). Interestingly, although FUBP1 primarily targets protein-coding genes (Fig. S3C), it also binds to the heavily-modified lncRNAs *MALAT1* and *NEAT1* in multiple clusters (Fig. S5A). Together, these findings demonstrate that FUBP1 preferentially binds to N_m_ sites in both caRNA and mRNA, consolidating its role as an N_m_-binding protein.

**Fig. 5.**
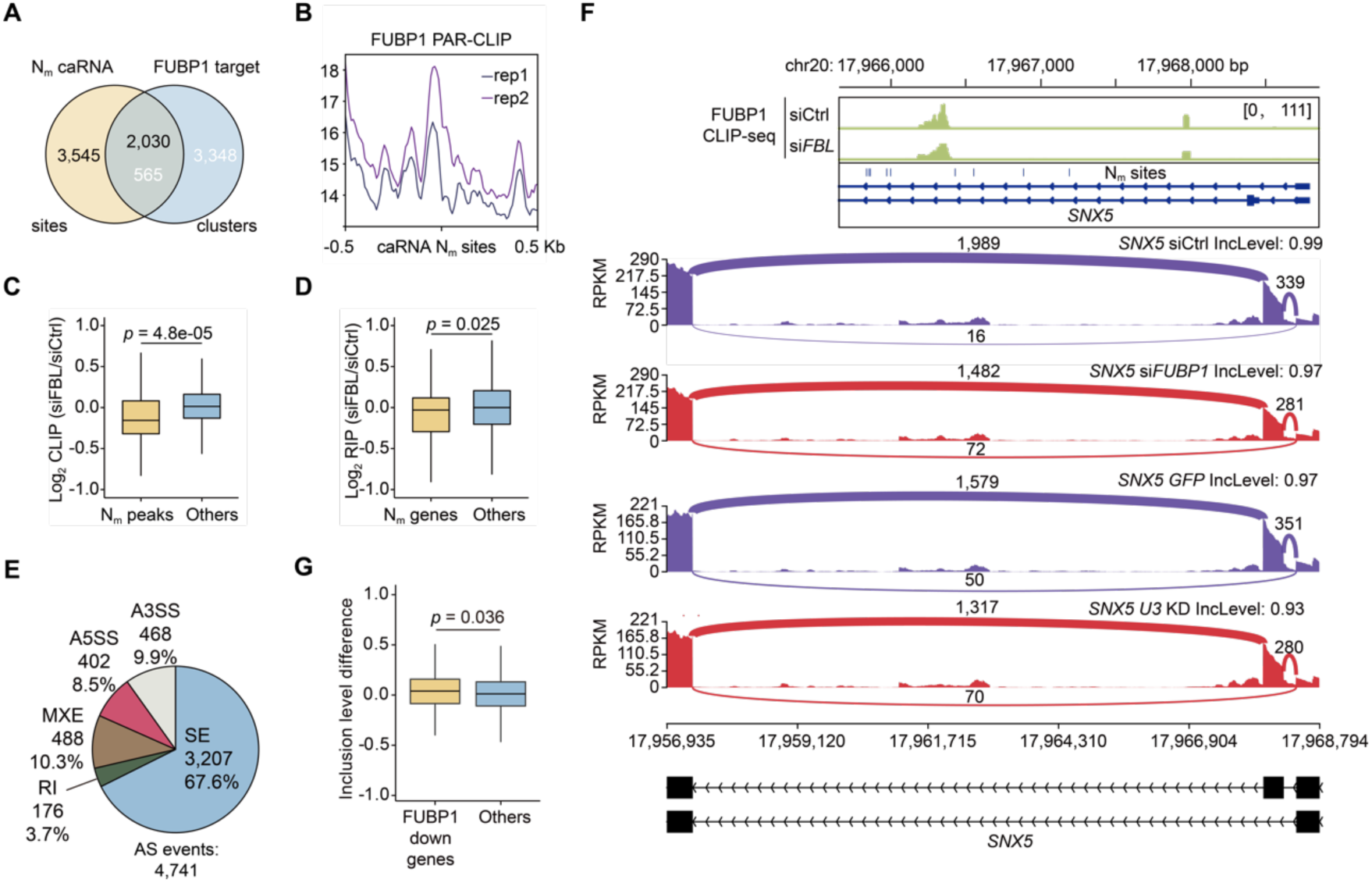
FUBP1 binds to N_m_-modified caRNA to affect splicing. (**A**) Overlap between caRNA N_m_ sites and FUBP1 PAR-CLIP clusters extending 200 bp on each side. (**B**) Metagene plot of FUBP1 PAR-CLIP signals at HepG2 caRNA N_m_ sites. (**C**) FUBP1 CLIP signal changes at peaks overlapping with N_m_ sites (N_m_ peaks) versus others after *FBL* KD. (**D**) FUBP1 RIP signal changes at gene transcripts with N_m_ modifications (N_m_ genes) versus others after *FBL* KD. (**E**) Profile of the FUBP1-dependent AS events. (**F**) Representative IGV tracks of differential FUBP1 CLIP-seq after si*FBL* and differential SE following si*FUBP1* or *U3* KD. (**G**) Inclusion level differences of SE events after FUBP1 depletion in genes with downregulated FUBP1 CLIP-seq peaks after si*FBL* (FUBP1 down genes) versus other genes (Others).

If FUBP1 indeed acts as a binding protein for N_m_, its binding should be affected by the depletion of N_m_. To test this, we conducted *FBL* KD. As PAR-CLIP provides mutation-based identification with less reliable measurement of differential peaks, we performed Crosslinking Immunoprecipitation (CLIP-seq) of FUBP1 following FBL depletion.

Consistent with our PAR-CLIP results, FUBP1 CLIP-seq peaks were predominantly located within intronic regions and protein-coding genes (Fig. S5B-C). Differential peak analysis revealed a decrease in FUBP1 peak signals overlapping with N_m_ sites, while the overall peak intensity remained largely unchanged (Fig. 5C). Similarly, gene-based differential binding measured by FUBP1 RNA Immunoprecipitation sequencing (RIP-seq) also demonstrated a significant decrease of N_m_-modified genes compared to unmodified ones after *FBL* KD (Fig. 5D). These findings collectively confirmed that FUBP1 binding to N_m_-modified RNA is indeed affected by the presence of N_m_ modification.

After confirming FUBP1 as a caRNA N_m_-binding protein, we aimed to explain the N_m_-dependent splicing regulation by FUBP1. FUBP1 facilitates bridging of 5′ and 3′ splice sites(*31*), and its depletion primarily induces exon skipping(*28*). Consistent with this, we knocked down *FUBP1* in HepG2 cells (Fig. S5D) and observed that SE was the main AS event (Fig. 5E). This aligns with the N_m_-dependent splicing regulation observed after *FBL* and *U3* KD (Fig. 4G and S4D). Among the SE events, upregulation occurred more frequently than downregulation (Fig. S5E), consistent with the biased upregulation induced by FBL or *U3* depletion (Fig. S4E). To further elucidate the relationship between N_m_ modification, FUBP1 binding, and splicing changes, we examined SE events in genes with caRNA N_m_ modifications. We found that N_m_-modified genes showed greater upregulation of SE events after FUBP1 depletion than non-modified genes, supporting that FUBP1 mediates N_m_-dependent inhibition of exon skipping. (Fig. 5F, S5F and G).

Similarly, genes with reduced FUBP1 binding following FBL depletion exhibited higher SE upregulation after *FUBP1* KD (Fig. 5G), further reinforcing the causal connection between N_m_ modification, FUBP1 binding, and splicing changes.

## Discussion

N_m_ modifications impact the physical properties of mRNA, exerting crucial regulation on the translation efficiency of modified genes(*18*). However, the “reader” proteins of N_m_ modifications on protein-coding genes remain elusive. Characterization of N_m_-binding proteins could reveal N_m_-dependent function. In this study, we employed RNA affinity purification followed by LC-MS/MS to identify candidate N_m_-binding proteins.

Immunoblotting validated the enrichment of FUBP1, KHSRP, FUBP3, IGF2BP1, IGF2BP3, FMR1, and NCOA5. Published eCLIP datasets of KHSRP, FUBP3, IGF2BP1, and IGF2BP3 further supported their enrichment around endogenous mRNA N_m_ sites.

While functions of IGF2BP proteins aligns with the effect of mRNA N_m_ on RNA expression levels, the identified FUBP family proteins and NCOA5 suggest unrecognized realms of nuclear N_m_ regulation.

We focused on FUBP1 and validated its role as an N_m_-binding protein. EMSA confirmed the direct interaction between FUBP1 and N_m_-modified RNA, while PAR-CLIP confirmed enrichment of FUBP1 around internal mRNA N_m_ sites. To further examine nuclear functions of N_m_ mediated through FUBP1, we profiled N_m_ modification site on caRNA by N_m_-mut-seq. We identified 5,575 caRNA N_m_ sites, with more than half of caRNA N_m_ sites in protein-coding genes localized in introns. We found that intronic N_m_ displayed enrichment around 5′ and 3′ splice sites, suggesting N_m_-mediated splicing regulation. Previous snoKARR-seq data revealed that *U3* is the predominant snoRNA responsible for N_m_ installation on caRNA. Depletion of *U3*, as well as methyltransferase FBL, led to upregulation of SE events, supporting a role of N_m_ in splicing regulation.

Interestingly, 2,228 identified N_m_ sites were densely populated on 323 protein-coding genes, suggesting cooperation among N_m_ sites in recruiting binding proteins. This may compensate for the modest preference of FUBP1 for a single N_m_ modification observed in EMSA experiments, contributing to significant enrichment of FUBP1 at N_m_-modified regions.

With the profiled N_m_ sites on caRNA, we validated the overlap of FUBP1 binding with caRNA N_m_ sites. To further explore the causal relationship, we disrupted N_m_ by FBL depletion, and detected impaired FUBP1 binding at N_m_-modified regions by CLIP-seq and RIP-seq. This further supports FUBP1 as the caRNA N_m_-binding protein. FUBP1 depletion upregulates SE events preferentially in N_m_-modified genes and genes with downregulated FUBP1 binding after FBL depletion, confirming its role in mediated N_m_-dependent splicing regulation. Overall, our findings identify FUBP1 as a caRNA N_m_-binding protein and uncover a new function of N_m_ in splicing modulcation through FUBP1.

## Materials and Methods

### Cell Culture

HepG2 cells (HB-8065) were cultured with media containing Dulbecco’s modified Eagle’s medium (Gibco, 11995040), 10% fetal bovine serum (FBS) (Gibco, 2614079) and 1% penicillin–streptomycin (Gibco, 15140122), at 37°C with 5% CO_2_ in the environment. Cells were passaged when reaching ∼90% confluency at 1:4 ratio. Mycoplasma were tested by PCR with primers gggagcaaacaggattagataccct and tgcaccatctgtcactctgttaacctc every half a year.

In the siRNA-mediated knockdown assay, cells at ∼90% confluency were passaged at 1:4 ratio into 15 cm plates. Within 16 hours after passaging, 60 μL Lipofectamine™ RNAiMAX Transfection Reagent (Invitrogen, 13778150) and 200 pmol siRNA were diluted in 1mL Opti-MEM™ I Reduced Serum Medium (Gibco, 31985070), respectively. The solutions were mixed together and incubated at room temperature for 5 mins before added into the cell culture. Knockdown reactions for fewer cells were scaled down based on the bottom area of culture plates. The siRNA used in this paper include siControl (Qiagen, 1027310), si*FBL* (Qiagen, SI04164951), siControl2 (Invitrogen, 4390846) and si*FUBP1*(Invitrogen, s16966). Cells were harvested 48 hours post transfection. KD efficiency was measured by qPCR with primers from Origene (HP205317).

*U3* KD was conducted with *U3* ASO

(mU*mC*mA*mC*mC*T*T*C*A*C*C*C*T*C*T*mC*mC*mA*mC*mU) controlled by GFP ASO

(mC*mU*mG*mC*mC*A*T*C*C*A*G*A*T*C*G*mU*mU*mA*mU*mC).

Transfection was done by Lipofectamine™ 3000 transfection reagents (Invitrogen, L3000015), where 600 pmol ASO and 8 μL P3000™ reagents were added to 125 μL Opti-MEM™ medium. In parallel, 6 μL Lipofectamine™ 3000 reagents were diluted in 125 μL Opti-MEM™ medium. The two mixtures were mixed together and incubated at room temperature for 10 min, and then added to 2 mL media in a 6-well plate well. HepG2 cells corresponding to 1/3 of a 10 cm plate at ∼90% confluency were then added to the transfected media. Cell were cultured for 72 hours before harvest. KD efficiency was measured by qPCR primer pair CGTGTAGAGCACCGAAAACC and CACTCAGACCGCGTTCTCTC.

### Immunoblotting

Samples were lysed in 2X NuPAGE™ LDS sample buffer (Invitrogen NP0007) supplemented with 1:20 (v/v) 2-Mercaptoethanol (Sigma-Aldrich, M6250-1L). After incubation at 95°C for 10 mins, the denatured samples were loaded into 4-12% NuPAGE Bis-Tris gels (Invitrogen, NP0322BOX) and transferred onto nitrocellulose membranes (Bio-rad, 1620115). Samples were blocked by 5% BSA (Fisher Scientific, BP1600-1) in PBST (Thermo Scientific, 28352) for 30 mins, followed by overnight incubation at 4°C in primary antibodies with designated dilution ratios in 3% BSA diluted by PBST. Membranes were washed 3 times and incubated in the secondary antibody conjugated to HRP at room temperature for 1 hour. Protein signals were developed by SuperSignal™ West Dura Extended Duration Substrate (Thermo Scientific, 34075). Antibodies used in this study and their dilution ratios are: Anti-FUBP1 (abcam, ab192867), 1:1000; Anti-KHSRP (CST, 13398S), 1:1000; Anti-FUBP3 (abcam, ab181025), 1:1000; Anti-IGF2BP1 (MBL international, RN007P), 1:1000; Anti-IGF2BP3 (MBL international, RN009P), 1:1000; Anti-NCOA5 (Proteintech, 20175-1-AP), 1:500; Anti-FMP1 (abcam, ab17722), 1:1000; Anti-rabbit IgG linked with HRP (CST, 7074S), 1:2000.

### RNA affinity purification

The experiment followed the protocol in the previous publication(*45*) with adjustment. 30 μL Dynabeads™ MyOne™ Streptavidin C1 beads suspension was washed with RNA binding buffer (50 mM HEPES-HCl pH 7.5, 150 mM NaCl, 0.5% NP-40 substitute, 10 mM MgCl2) and incubated in RNA binding buffer supplemented with 100 ug/mL yeast tRNA (Invitrogen, AM7119) for 1 hour at 4°C with rotation. After two rounds of washing, 400 pmol N_m_-modified or control probes were incubated with beads suspended in RNA binding buffer for 30 mins at 4°C with rotation. Beads conjugated with oligos were washed with RNA wash buffer (50 mM HEPES-HCl pH 7.5, 250 mM NaCl, 0.5% NP-40 substitute and 10 mM MgCl2) and then with protein incubation buffer (10 mM Tris-HCl pH 7.5, 150 mM KCl, 1.5 mM MgCl2, 0.1% NP-40 substitute and 0.5 mM DTT) twice.

HepG2 cell pellets in the volume of 45 uL were lysed in 400 ul lysis buffer (50 mM Tris pH 7.5, 100 mM NaCl, 1% NP-40 substitute, 0.5% sodium deoxycholate, 100 × protein inhibitor cocktail (Sigma-Aldrich, P8340) and 100 × SUPERase•In™ RNase Inhibitor (Invitrogen, AM2696)) for 30 mins at 4°C with rotation. The supernatant of the lysate was harvested by centrifugation at 12,000g for 15 mins, and separated to two equal halves after saving 5% as input. The lysate was incubated with beads conjugated with oligos, supplemented with 50 ug/mL tRNA, 0.5 mM DTT and 100 × SUPERase•In, incubated at room temperature for 30 mins and 4°C for 90 mins with rotation. The beads were washed with protein incubation buffer for 3 times before removal of all supernatant.

### LC-MS/MS analysis

Samples were prepared with 3 replicates, harvested on dry beads and frozen in dry ice when shipped to the MS center. The beads were heated in 3x reducing LDS sample buffer with 15 mM DTT and 2 M biotin for 10 mins at 95°C, and the supernatant was loaded on 12% Bis-Tris propane SDS-PAGE gel for removal of detergent. The gel was run shortly and stained with colloidal coomassie blue for gel cut of the whole lane. Gel pieces were reduced with DTT, alkylated with iodoacetamide, washed properly and digested with trypsin overnight at 37°C. The extracted peptides were dried down and re-dissolved in 2.5% acetonitrile-water solution with 0.1% formic acid, then run by nanoLC-MS/MS using a 2-hour gradient on a 0.075mmx250mm C18 column feeding into an Orbitrap Eclipse mass spectrometer.

The quantitation was done by Proteome Discoverer (Thermo; version 2.4). All MS/MS samples were searched with Mascot (Matrix Science, London, UK; version 2.6.2) utilizing cRAP_20150130.fasta (124 entries); uniprot-human_20201207 database (75777 entries) assuming trypsin digestion, with a fragment ion mass tolerance of 0.02 Da and a parent ion tolerance of 10.0 PPM. The specified variable modifications included asparagine and glutamine deamidation, methionine oxidation, lysine and arginine methylation and cysteine carbamidomethylation. Peptides were validated by Percolator with a 0.01 posterior error probability (PEP) threshold. The data were searched with a decoy database to set the false discovery rate to 1%. The peptides were quantified using the precursor abundance based on intensity, with the peak abundance normalized by total peptide amount. The sum of peptide group abundances for each sample were normalized to the maximum sum of the analyzed samples. The protein ratios were calculated using summed abundance for each replicate separately and the geometric median of the resulting ratios was used as the protein ratios. To compensate for missing values in some of the replicates, the replicate-based resampling imputation mode was used. The significance of differential expression was generated by ANOVA test and the *p*-values were adjusted by the Benjamini-Hochberg method.

### Protein purification

FUBP1-strep were expressed in 100 ml Expi293F cells for 48 hours, lysed in lysis buffer (20 mM Tris pH 8.0, 150 mM KCl, 2 mM EDTA, 100 × PMSF, 1% NP-40 substitute and 0.5 mM DTT) at 4°C for 30 mins. Supernatant was harvested by centrifugation, and diluted to 3-fold volume before passing through 0.22 μm filter. Samples were loaded to 200 μl Strep-Tactin® Sepharose® resin (IBA, 21201010) and incubated at 4°C for 1 hr. The resin was flowed through by lysate twice and washed with 25 mL wash buffer (50 mM Tris pH 8.0, 250 mM NaCl, 0.05% NP-40 substitute, 0.5 mM DTT and 100 × PMSF). Protein was eluted by elution buffer (50 mM Tris, pH 8.0, 250 mM NaCl, 0.01% NP-40 substitute, 0.5 mM DTT and 2.5 mM dethiobiotin) × 6 with 100 uL each time, concentrated with 30 kD Amicon® Ultra Centrifugal Filter (Millipore, UFC503008) and exchanged to storage buffer (50 mM Tris, pH 8.0, 150 mM KCl, 0.1 mM EDTA and 20% Glycerol). The purified protein was aliquoted, snap frozen in liquid nitrogen and stored at -80°C.

### EMSA

Probes were refolded by incubation at 75°C for 2 mins and natural cooling to room temperature for 10 mins. FUBP1 proteins of designated concentrations were incubated with 20 nM FAM-labeled oligos in binding buffer (10mM Tris pH 7.5, 140 mM KCl, 10mM NaCl, 1 mM MgCl2, 10% glycerol, 1 mM DTT and 1U/uL SUPERase•In™ RNase Inhibitor) at room temperature for 30 mins. The mixtures were loaded to 4-20% Novex™ TBE gels (EC62255BOX) that have been rerun at 90V for 30 mins at 4°C in 0.5 × TBE.

Gels were run for 2 hrs before imaging. The fluorescence signal intensity was quantified by imageJ, and Kd were calculated by GraphPad.

### UHPLC-QQQ-MS/MS quantification

HepG2 mRNA were purified by two rounds of polyA selection following the commercial protocol of Dynabeads™ mRNA DIRECT™ Purification Kit (Invitrogen, 61011). HepG2 chromatin were purified following the published protocol reported previously(*7*), with adjustment of lysis buffer to 10 mM Tris-HCl pH 7.5, 0.15% NP-40 substitute and 150 mM NaCl. The caRNA were purified from fractionated chromatin, followed by two rounds of ribominus reactions according to the commercial instruction of RiboMinus™ Eukaryote System v2 (Invitrogen, A15026). For both samples, 100 ng RNA was digested with Nuclease P1 (NEB, M0660S) at 37°C overnight, followed by digestion with Shrimp Alkaline Phosphatase (rSAP, NEB, M0371S) in rCutSmart™ buffer. The samples were diluted and filtered by 0.22 μm PVDF filter (Millipore, SLGVR33RB), then injected into a C18 reverse phase column coupled online to Agilent 6460 LC-MS/MS spectrometer in positive electrospray ionization mode. Nucleosides were quantified using nucleoside-to-base transitions of A_m_ (282 to 136), A (268 to 136), G_m_ (198.1 to 152.1), G (284 to 152), C_m_ (258.2 to 112), C (244 to 112). The signal intensity of N_m_ nucleotides was normalized to the corresponding unmodified nucleotides to enable comparison of samples with different length.

### PAR-CLIP

The experiment was performed with adapted protocol from previous reports(*46*). Two replicates of 150 million HepG2 cells were cultured with 200 μM 4SU for 14 hrs. Cells were crosslinked by 365 nm UV at 1500 J/m^2^ twice, harvested and lysed by iCLIP lysis buffer (50 mM Tris pH 7.5, 100 mM NaCl, 1% NP-40 substitute, 0.1% SDS, 0.5% sodium deoxycholate, 100 × protein inhibitor cocktail and 100 × SUPERase•In™ RNase Inhibitor) at 4°C for 15 mins with rotation. To release FUBP1 proteins associated with the chromatin, cell lysate was supplemented with 1% SDS, sonicated at 30% amplitude with 2s:4s cycles for 1 min on ice. The lysate was 10-fold diluted by iCLIP lysis buffer without SDS, centrifuged to save the supernatant, and treated with 0.2 U/μL RNase T1 (Thermo, EN0642) at 22°C for 15 mins before quenched on ice for 5 mins. Protein G beads (Invitrogen, 10009D) were conjugated with 20 μg FUBP1 antibody (abcam, ab192867) by incubation at 4°C for 1hr, then washed and mixed with RNase T1-treated lysate to rotate at 4°C for 4 hrs. Beads were washed for three times with CLIP wash buffer (50mM Tris pH 7.5, 300 mM KCl, 0.05% NP-40 substitute, 1000 × protein inhibitor cocktail and 1000 × SUPERase•In™ RNase Inhibitor), digested by 10 U/μL RNase T1 at 22°C for 8 mins, before they were washed by CLIP High salt buffer (50mM Tris pH 7.5, 500 mM KCl, 0.05% NP-40 substitute, 1000 × protein inhibitor cocktail and 1000 × SUPERase•In™ RNase Inhibitor) and PNK buffer without DTT for 3 times each and underwent end repair by T4 PNK (NEB, M0201L). The immunoprecipitation was validated by biotinylation, eluted by proteinase K digestion (Thermo Scientific, EO0491), with purified RNA constructed to libraries by NEBNext^®^ Small RNA Library Prep Set (NEB, E7330S). Sequencing was performed by Illumina NovaSeq6000 reading pair-end 50 bp.

### Bioinformatic analysis of PAR-CLIP

Adapters were trimmed by cutadapt(*47*), and reads were mapped to hg38 human genome by HISAT2(*48*), with parameter “--reorder --no-unal --pen-noncansplice 12”. Duplicates were filtered by Picard MarkDuplicates. R1 reads with mapping quality higher than 30 were used for identification of FUBP1 binding clusters by wavClusteR(*49, 50*), with removal of reads containing “^” in the MD tag. The identified clusters were overlapped, then annotated by Homer(*51*) annotatepeaks. Metaplots centering at N_m_ sites were generated by DeepTools(*52*). Motifs were identified by Homer findMotifsGenome with parameter “-rna -size 200 -len 6”.

### CLIP-seq

The experiment followed the procedure in previous publication(*53*). Three replicates of 150 million HepG2 cells were harvested after 48 hr of FBL knockdown, with the knockdown efficiency was validated by RT-qPCR with primers from Origene (HP205317). Samples were crosslinked by 1500 J/m^2^ UV at 254 nm for three times, lysed, centrifuged and digested with the same condition as PAR-CLIP. After saving 2% of the lysate as input, lysates were incubated with beads treated the same as PAR-CLIP. The following steps were exactly the same, with the exception that input samples were treated separately with 10 U/μL RNase T1 at 22°C for 8 mins, followed by proteinase K digestion and end repair in the next. Sequencing by Illumina NovaSeq X was conducted for single-end 100 bp.

### Bioinformatic analysis of CLIP-seq

Cutadapt(*47*) was used to trim the adapters, and HISAT2(*48*) mapping to the reference genome (hg38) was performed, with parameter “--reorder --no-unal --pen-noncansplice 12”. Peaks were identified by Piranha(*54*) with input samples as the covariate, and bin size designated as 50 bp. The identified peaks of siControl and siFBL were merged and annotated by Homer(*51*) annotatepeaks. Differential binding was analyzed by DiffBind, with relative log expression normalization considering the background (normalize = DBA_NORM_RLE, background = TRUE). Downregulated peaks were defined as log_2_ fold change < -0.58 and FDR < 0.1. Peaks within 2 kb of N_m_ sites were considered as N_m_ modified peaks for computing binding fold change.

### RNA-seq

HepG2 cell RNA was harvested by TRIzol™ reagent (Invitrogen, 15596026) after 48 hours of siRNA-mediated KD or 72 hours of ASO-mediated KD. RNA was purified with manufacturer’s procedures, and the KD was measured by RT-qPCR with primers from Origene (HP207343). After polyA RNA selection with Dynabeads™ mRNA DIRECT™ Purification Kit and the samples were constructed to libraries with SMARTer® Stranded Total RNA-Seq Kit v2 - Pico Input Mammalian (Takara, 634412). Next generation sequencing was performed with Illumina NovaSeq6000 reading pair-end 150 bp.

### RIP-seq

Samples were harvested after 48 hours of *FBL* KD. The KD efficiency was validated by RT-qPCR. Cells were lysed by iCLIP lysis buffer at 4°C for 15 mins with rotation, followed by centrifugation at 13,000 g for 15 mins. Protein G beads were conjugated with 16 μg FUBP1 antibody by incubation at 4°C for 1 hour, then washed and mixed with lysate supernatant to rotate at 4°C for 4 hours. Beads were washed with CLIP wash buffer for three times and eluted by proteinase K digestion (Thermo Scientific, EO0491) Input samples harvested before immunoprecipitation was also digested with proteinase K for RNA recovery. The purified RNA was constructed for library with SMARTer® Stranded Total RNA-Seq Kit v2 - Pico Input Mammalian. Next generation sequencing was performed with Illumina NovaSeq6000 reading pair-end 150 bp.

### Bioinformatic analysis for RNA-seq and RIP-seq

Adapters were trimmed by cutadapt(*47*), and reads were mapped to hg38 human genome by HISAT2(*48*), with parameter “--reorder --no-unal --pen-noncansplice 12”. RIP-seq reads were count for each gene by HTseq(*55*), and differential expression analysis by DESeq2(*56*) was conducted on IP versus input samples, where the fold change was considered as enrichment of each gene. The enrichment was further compared between control and FBL knockdown samples to measure the differential binding of FUBP1 upon Nm depletion.

The AS events were detected by rMATS(*57*), where we utilized the input samples of RIP-seq for analysis upon *FBL* KD. Only junction reads were used for calculation of AS scores. Events with FDR<0.1 were considered as AS events.

### Nm-mut-seq

HepG2 caRNA was harvested in the same procedures as UHPLC-QQQ-MS/MS analysis. rRNA was removed by RiboMinus™ Eukaryote System v2, and the residual RNA was cleaned up by RNA Clean & Concentrator™-5 (Zymo Research, R1014) with removal of short RNA < 200 nt. The construction of Nm-mut-seq followed the published procedures(*27*), using 3′ linker 5’rApp- NNNNNATCACGAGATCGGAAGAGCACACGTCT-3SpC3 and 5′ SR adapter supplied in the NEBNext^®^ Small RNA Library Prep Set. Sequencing by Illumina NovaSeq X was conducted with single-end 100 bp.

### Bioinformatic analysis for Nm-mut-seq

Adapters were trimmed by cutadapt(*47*), and duplicates marked by UMI was removed using BBMap61. Reads were mapped to hg38 human genome by TopHat2(*58*), with parameters “-g 4 -N 3 --read-edit-dist 3”. Mutations to T were identified by JACUSA(*40*), with further selection of mutated sites in all three replicates having sequencing depth ≥ 10, mutated read depth ≥ 3, mutation ratios > 3-fold of input mutation ratios, and mutation ratios > 1.5-fold of background mutation ratios. Nm sites were annotated by Homer(*51*) annotatepeaks.

### Statistical Analysis

*P*-values annotated in figures were quantified based on two-tailed student t-test. Chi-squared test was used to statistically test the preferential occurrence of different splicing events with increased or decreased inclusion level differences. The Kd value and SD for EMSA was fitted by prism under the mode of “one site – specific binding with Hill slope”.

## Acknowledgments

We are grateful to the Genomics Facility at University of Chicago for high-throughput sequencing, the Mass Spectrometry Facility at University of Chicago for UHPLC-QQQ-MS/MS, and the Proteomics & Metabolomics Facility at University of Nebraska - Lincoln for LC-MS/MS analysis.

## Author contributions

Conceptualization: CH. Methodology: CH, BG, BJ, LZ, BL, WL, LC. Investigation: Ch, BG. Visualization: CH, BG, ZZ. Supervision: CH. Writing— original draft: CH, BG. Writing—review and editing: CH, BG, ZZ.

## Data and materials availability

All data are available in the main text and/or the Supplementary Materials. The sequencing data are accessible in the Gene Expression Omnibus through accession number GSE294792, GSE294793, and GSE294794.

ENCODE datasets of eCLIP experiments targeting KHSRP (ENCSR366DGX), FUBP3 (ENCSR486YGP), IGF2BP1 (ENCSR744GEU) and IGF2BP3 (ENCSR993OLA) were used for analysis.

---

**Fig. S1.**
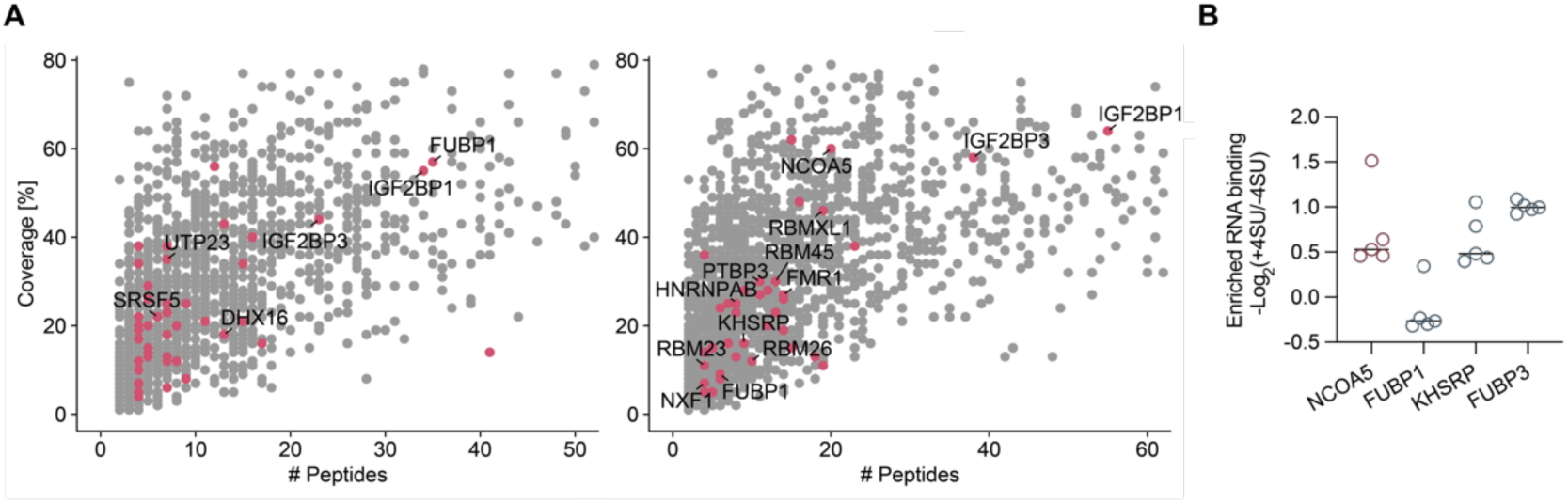
Identification of candidate N_m_-binding proteins. (**A**) Peptides from proteins that showed binding to RNA Gm-1 (left) and Am-2 (right) probe. N_m_-binding protein candidates (red) should have log_2_ fold change > 1.58, adjusted *p*-value < 0.1 and peptide number ≥ 4. # Peptides: peptide number. Coverage: peptide coverage of the protein. (**B**) Enriched RNA-binding of 5 most highly enriched peptides bound to RNA represented by -log_2_ fold change of +4SU/-4SU samples in published RBR-ID of K562 cells. Peptides were filtered for adjusted *p*-value < 0.05 and ranked by with fold change.

**Fig. S2.**
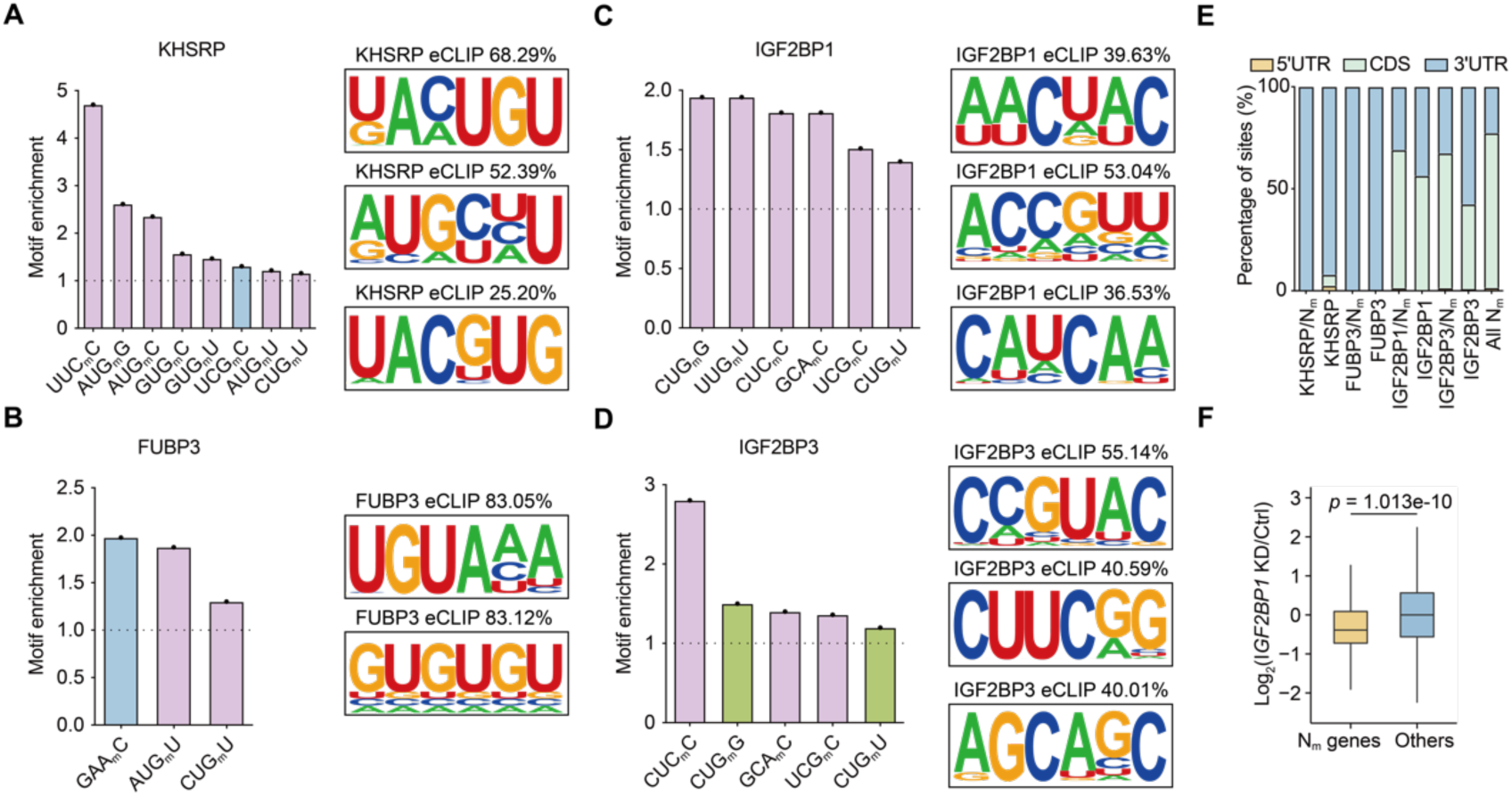
The target properties of N_m_-binding proteins. (**A-D**) Enrichment of N_m_ motifs at the N_m_-modified binding sites (left) and the enriched motifs across all binding sites from ENCODE eCLIP dataset (right) of KHSRP (A), FUBP3 (B), IGF2BP1 (C), and IGF2BP3 (D). N_m_ motifs resembling the enriched RNA binding motifs of the respective RBPs are shown in pink. N_m_ motifs that are also enriched by other members of the protein families (KHSRP /FUBP3 or IGF2BP1/IGF2BP3) are colored in green. (**E**) Percentage of N_m_ sites bound by each N_m_-binding protein (N_m_-binding protein/N_m_), total binding peaks of each N_m_-binding protein, or all N_m_ sites across different mRNA regions. (**F**) Fold changes in RNA expression following *IGF2BP1* KD for genes with confident N_m_ modification (N_m_ genes) versus other genes (Others).

**Fig. S3.**
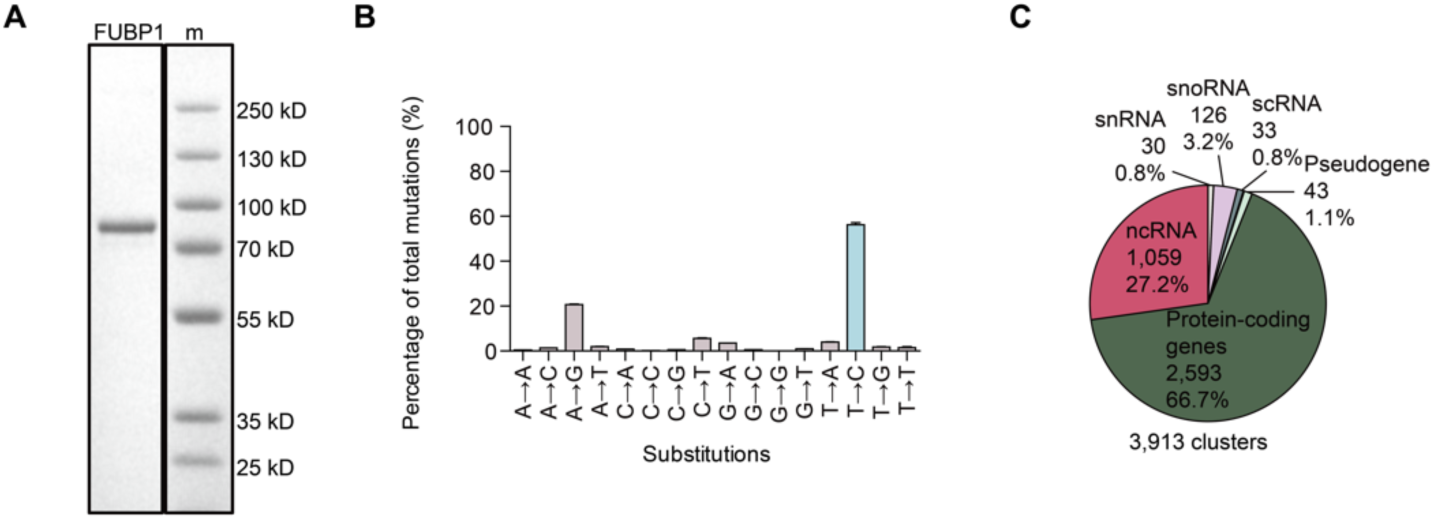
Analysis of FUBP1 binding profile. (**A**) Coomassie staining of purified FUBP1-strep expressed in Expi293 cells. (**B**) Base conversion ratios in FUBP1 PAR-CLIP. (**C**) Distribution of FUBP1 PAR-CLIP clusters across different gene types.

**Fig. S4.**
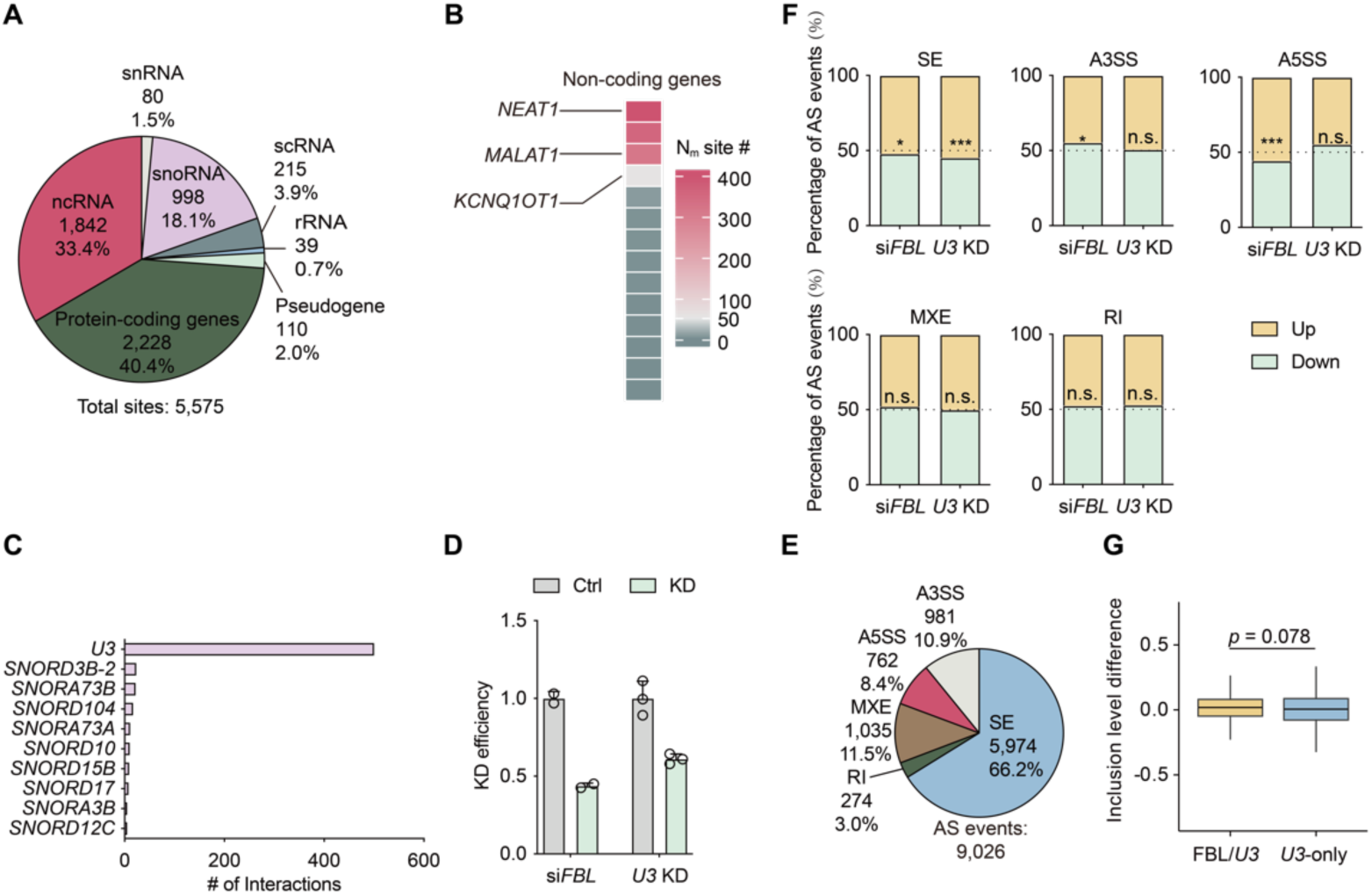
caRNA N_m_ distribution and its implied function in splicing regulation. (**A**) Distribution of 5,575 caRNA N_m_ sites across different gene types. (**B**) Heatmap showing numbers of N_m_ sites in various non-coding caRNAs. (**C**) Number of snoRNA-caRNA interactions detected by snoKARR-seq at N_m_ sites within protein-coding genes. (**D**) KD efficiency measured by qPCR normalized to *ACTB*. (**E**) Profile of *U3*-dependent AS events identified with FDR < 0.1. (**F**) Percentage of increased (Up) or decreased (Down) AS events after FBL depletion or *U3* KD. *, *p*-value < 0.05; ***, *p*-value < 0.005; n.s., not significant. (**G**) Inclusion level differences of FBL/*U3* or *U3*-only SE events.

**Fig. S5.**
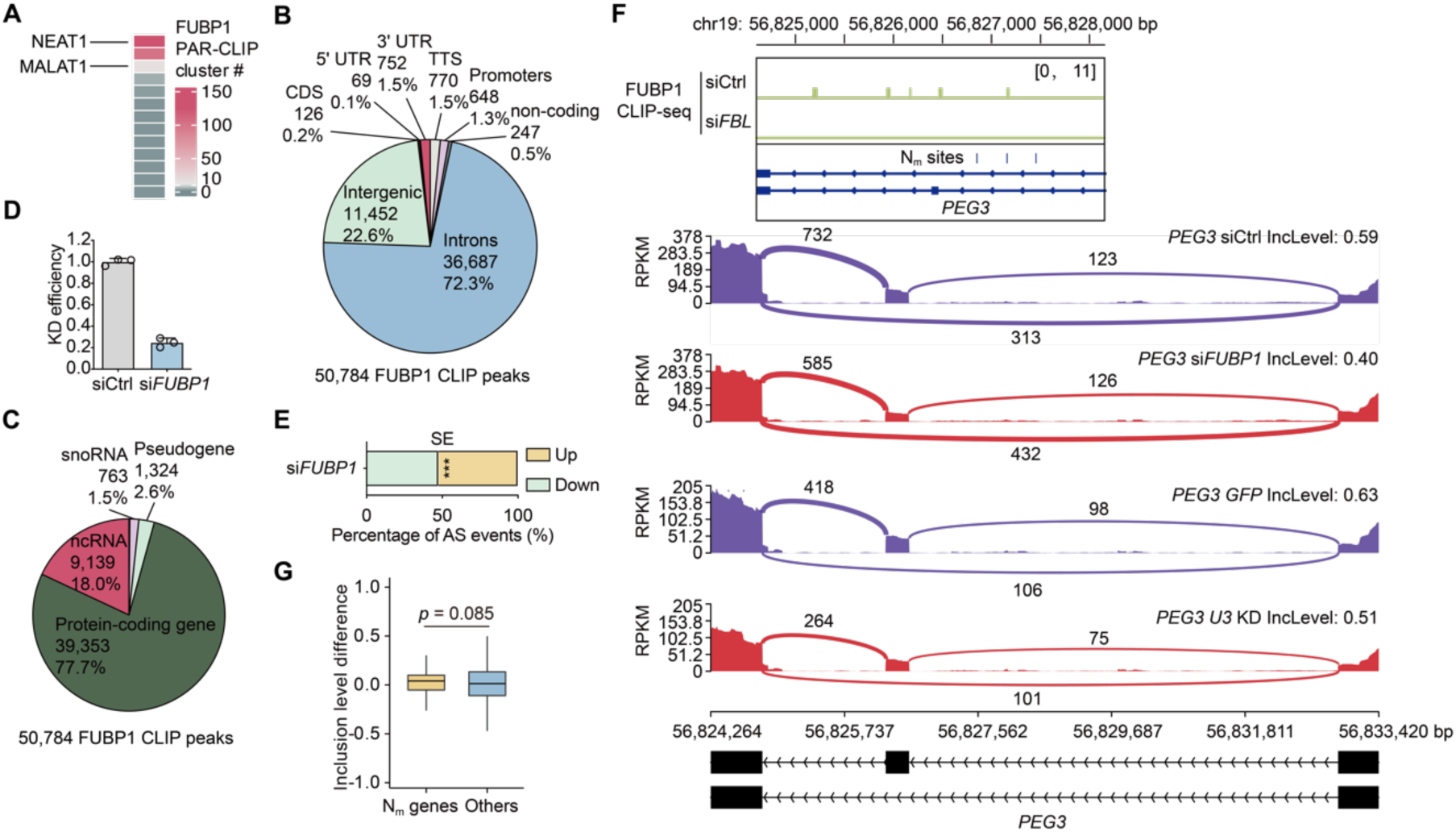
FUBP1 binds to N_m_-modified caRNA to affect splicing. (**A**) Heatmap showing numbers of FUBP1 PAR-CLIP clusters in various non-coding RNA. (**B**) Distribution of FUBP1 CLIP-seq peaks across transcript elements. (**C**) Distribution of FUBP1 CLIP-seq peaks across gene types. (**D**) KD efficiency measured by qPCR normalized to *ACTB*. (**E**) Percentage of increased (Up) or decreased (Down) SE events after FUBP1 depletion. ***, *p*-value < 0.005. (**F**) Representative IGV tracks of differential FUBP1 CLIP-seq after si*FBL* and differential SE following si*FUBP1* or *U3* KD. (**G**) Inclusion level differences of SE events in genes with caRNA N_m_ sites (N_m_ genes) versus other genes (Others) after FUBP1 depletion.

## Supplementary Tables S1-9

Table S1. Protein enrichment from Gm-1/Ctrl-1 oligo pull down followed by proteomics.

Table S2. Protein enrichment from Am-2/Ctrl-2 oligo pull down followed by proteomics.

Table S3. Annotation of FUBP1 PAR-CLIP clusters.

Table S4. Gene annotation of caRNA Nm sites identified by Nm-mut-seq.

Table S5. Skip exons of FBL-depleted HepG2 cells.

Table S6. Skip exons of *U3*-depleted HepG2 cells.

Table S7. DiffBind analysis of FUBP1 CLIP-seq peaks with FBL depletion.

Table S8. DESeq2 analysis of FUBP1 RIP-seq with FBL depletion.

Table S9. Skip exons of FUBP1-depleted HepG2 cells.

